# 14-3-3 protein Bmh1 triggers short-range compaction of mitotic chromosomes by recruiting sirtuin deacetylase Hst2

**DOI:** 10.1101/2020.06.09.142752

**Authors:** Neha Jain, Petra Janning, Heinz Neumann

## Abstract

Mitotic chromosome compaction is licensed by kinetochores in yeast. Recruitment of Aurora kinase B elicits a cascade of events starting with phosphorylation of H3 S10, which signals the recruitment of lysine deacetylase Hst2 and the removal of H4 K16ac. The unmasked H4 tails interact with the acidic patch of neighbouring nucleosomes to drive short-range compaction of chromatin. Here, we demonstrate that the interaction of Hst2 with H3 S10ph is mediated by 14-3-3 protein Bmh1. As a homodimer, Bmh1 binds simultaneously to H3 S10ph and the phosphorylated C-terminus of Hst2. The Hst2-Bmh1 interaction is cell cycle dependent, reaching its maximum in M phase. Furthermore, we show that phosphorylation of C-terminal residues of Hst2 stimulates its deacetylase activity. Hence, the data presented here identify Bmh1 as a key player in the mechanism of licensing of chromosome compaction in mitosis.

**Key Points:**

- 14-3-3 protein Bmh1 bridges the interaction of Hst2 with phosphorylated H3 tails
- Hst2 is multiply phosphorylated on its unstructured C-terminal tail
- The interaction of Bmh1 with Hst2 is cell cycle dependent
- Hst2 phosphorylation enhances its enzymatic activity

## Introduction

In mitosis, cells condense their chromosomes into compact, cylindrical bodies to ensure their faithful mechanical transport during cell division (1). Therefore, ring-shaped protein complexes, the condensins, compact mitotic chromosomes into helical arrays of chromatin loops (2). Condensins are believed to act by loop extrusion (3), fuelled by hydrolysis of ATP by their ATPase domains. Loop extrusion has been observed in single molecule experiments (4,5) and is supported by structural analysis of DNA-bound condensin subunits (6).

Depletion of condensins has dramatic consequences for mitotic chromosome architecture and mechanical stability (7-11). However, additional forces and factors must contribute to chromosome condensation because even in the absence of condensins chromatin still aggregates in mitosis (10,12). Among the many factors that may contribute to this phenomenon, posttranslational modifications (PTMs) of histones are particularly attractive (13,14). Histone PTMs may contribute to chromosome condensation in various ways: they could signal the recruitment of effector proteins by serving as recognition marks, might influence the activity of enzymes involved in condensation or directly control an inherent tendency of chromatin to condense.

The most intensely studied mitotic PTM is phosphorylation of Histone H3 Ser-10 (15). This hallmark of mitotic chromatin is initially deposited at centromeres by Aurora Kinase B as part of the Chromosomal Passenger Complex (CPC) (16). In budding yeast, H3 S10ph signals the recruitment of lysine deacetylase (KDAC) Hst2 to chromatin in mitosis (17). Hst2 in turn removes acetyl groups from lysine-16 of nearby H4 tails, thereby enabling them to engage with the acidic patch of neighboring nucleosomes. This inter-nucleosomal interaction provides a driving force of chromatin compaction in mitosis, mediating hypercondensation of chromatid arms in late anaphase (17,18). This short-range compaction acts independently of the axial contraction by condensins (17,18). Mechanistically, compaction by inter-nucleosomal interactions is initiated by activation of Aurora kinase B at kinetochores. The signal subsequently propagates along chromosome arms in a shugoshin-dependent process, thereby coupling the ability of chromatin to condense to the presence of a centromere (19). This licensing of condensation by centromeres is a potential mechanism by which yeasts discriminate non-chromosomal DNA from chromosomes, protecting its progeny from infectious genetic material (19).

The spreading mechanism of short-range compaction is essential to create a chromosome-autonomous process. Little is presently known about the molecular mechanism of spreading. Here, we explore how Hst2 recognizes H3 S10ph and how this interaction is controlled by the cell cycle.

## Materials and Methods

### Plasmids and Strains

Overexpression of Hst2 proteins was from 2µ plasmids under the control of the HST2 promoter and terminator. For expression in *E. coli*, ORFs of HST2 and BMH1 were cloned in pCDF DUET with purification tags. Mutations were introduced by QuikChange mutagenesis. Details can be found in the Supplementary Materials. Yeast strains were constructed in the S288C background. Yeast strain were created according to standard procedures. The strains used in this study are listed in Table S1.

### Cell cycle arrests

To synchronize cells in G1, the mating pheromone α-factor (GenScript) was added to log phase cultures grown to OD_600_ of 0.4–0.6 (MATa yeast strain BY4741) to a final concentration of 15 μg/mL. Cells were incubated for 1 h at 30°C at which time an additional dose of 7.5 μg/mL α-factor was added and cells incubated for one more hour. Cells were monitored periodically by microscopy. Cells were collected by centrifugation at 4000 rpm for 2 min and processed for further experiments. To synchronize cells in metaphase, one dose of nocodazole (15 μg/mL, Sigma Aldrich) was added to log phase cultures (OD_600_=0.5) for 2 h. The cells were monitored by microscopy and collected as above. To synchronize cells in S phase, 100 mM hydroxyurea (Sigma Aldrich) were added to exponentially growing cells in YPD (OD_600_ ≈ 0.3) and incubated for two hours at 30°C, after which they were monitored by microscopy and collected as above.

### Protein purifications

#### Hst2

*Escherichia coli* BL21 (DE3) transformed with pCDF-His6-yHst2 were grown overnight at 37°C in LB containing 50 µg/mL spectinomycin. Next day, the preculture was used 1:50 to inoculate 4 L LB medium and protein expression induced with 0.5 mM IPTG at OD_600_ = 0.5, and incubation continued overnight at 15°C. Cells were harvested, suspended in 20 mM HEPES, pH 7.5, 200 mM NaCl, 20mM Imidazole, 3 mM β-mercaptoethanol and 1 mM PMSF supplemented with lysozyme (∼0.5 mg/mL), DNase (1 mg), protease inhibitor cocktail (Roche) and disrupted with a pneumatic cell disintegrator. Soluble His-yHst2 was purified by Ni-NTA affinity chromatography using 20 mM HEPES pH 7.5, 200 mM NaCl, 20 mM imidazole, 3 mM β-mercaptoethanol for binding and washing and additional 200 mM imidazole for elution. The eluate was concentrated and separated by gel filtration in 20 mM HEPES, pH 7.5, 100 mM NaCl, 0.5 mM TCEP on a 16/60 Superdex 75 column (GE healthcare, UK). Hst2-containing fractions were pooled, concentrated in a microfiltrator and stored in aliquots at –80°C.

#### Bmh1

Protein expression conditions were the same as for Hst2. Cells were lysed in 50 mM Tris pH 8, 300 mM NaCl, 3 mM β-mercaptoethanol, 10% glycerol. The supernatant was incubated with Pierce™ High Capacity Streptavidin Agarose, washed with the same buffer and Strep-Bmh1 eluted with additional 10 mM desthiobiotin. Strep-Bmh1 was further purified on a Superdex 200 column (GE Healthcare) in 20 mM Tris pH 8, 100 mM NaCl, 0.5 mM TCEP, 10% glycerol.

#### Hst2 S320ph and S324ph

BL21 ΔserB (DE3) cells containing pKW2-EF-Sep (a chloramphenicol resistant plasmid containing SepRS2, pSer-tRNAB4_CUA_ and EF-Sep (20)) were transformed with pCDF-His-yHst2 S320TAG or S324TAG. Cells were grown at 37°C in LB medium containing 50 µg/mL spectinomycin and 34 µg/mL chloramphenicol and used next day to inoculate 4 L LB-SC 1:50. Protein expression was induced at OD_600_=0.5 with 1 mM IPTG, and cells were harvested after 4 h at 37°C. Purification followed the same protocol as for unmodified Hst2. All proteins were stored at –80°C.

### Co-precipitation experiments

BY4741 cells were transformed with plasmid pRS423-Flag-Hst2 and grown to OD_600_=1.7–3.0. Cells from 1 L were resuspended in PBS supplemented with protease inhibitors (1 mM phenylmethanesulfonylfluoride and each 5 μg/mL chymostatin, leupeptin, aprotinin and pepstatin A) and (where indicated) Phosphatase inhibitor cocktail (PhosSTOP™ Sigma Aldrich). Subsequently, cells flash frozen cell nuggets were lysed by milling (RETSCH ZM 200 Ultra Centrifugal Mill), thawed and centrifuged (20000 rpm, 4°C for 15 minutes). The supernatant (total protein concentration ∼3 mg/ml) was incubated with ANTI-FLAG M2 agarose beads (Sigma Aldrich) at 4°C for 1 h with agitation. Beads were washed six times with 500 μL PBS containing 0.2% Triton^®^ X-100 and finally eluted by boiling in SDS sample buffer. Proteins were analyzed by SDS-PAGE and Western blot (antibodies are listed in Table S2).

### In vitro pulldowns

Purified, recombinant Bmh1 and Hst2 isoforms (0.3 mg each) were mixed with 20 μl 50:50 slurry of Avidin Agarose Resin (Fisher Scientific) in 80 µl of Bmh1 SEC buffer (20 mM Tris pH 8, 100 mM NaCl, 0.5 mM TCEP, 10% glycerol), incubated for 1 h at RT with shaking (300 rpm) and subsequently washed three times with PBS. Proteins were eluted by boiling in SDS-PAGE sample buffer, analyzed by 10% SDS-PAGE and stained with Instant Blue.

### Luciferase-based KDAC assay

KDAC activity was measured in continuous assay format using Firefly Luciferase (FLuc) K529ac (21). Reactions contained 200 nM Hst2 (unmodified or phosphorylated), NAD^+^ at concentrations from 0 – 4 mM, FLuc K529ac (suitably diluted to match the sensitivity of the luminometer) in 50 µl KDAC buffer (25 mM Tris/HCl pH 8.0, 137 mM NaCl, 2.7 mM KCl, 1 mM MgCl_2_, 1 mM GSH). To assay luciferase activity, an equal volume of 40 mM Tricine pH 7.8, 200 µM EDTA, 7.4 mM MgSO_4_, 2 mM NaHCO_3_, 34 mM DTT, 0.5 mM ATP and 0.5 mM luciferin was added and luminescence recorded for 30 min at 30°C in a FluoStar Omega Microplate Reader (BMG Labtech). All experiments were executed in triplicate, averaged and reactions without the enzyme were used for background subtraction. Initial rates were determined from the linear phase of the reactions. The kinetic parameters (apparent K_M_ and k_cat_) were obtained by fitting the data to the non-linear regression Michaelis-Menten model in GraphPad Prism 8 software.

## Results and Discussion

H3 peptides phosphorylated at Ser-10 efficiently recovered Hst2 from yeast whole-cell extracts, demonstrating the significance of H3 S10 phosphorylation in recruiting Hst2 to mitotic chromatin (17). However, recombinant Hst2 purified from *E. coli* did not interact with H3 S10D peptides (a mutation that efficiently mimics H3 S10ph *in vivo* (17,19,22)) (Supplementary Fig. S1), indicating that additional factors or PTMs on Hst2 are needed to mediate the interaction with H3 S10ph.

In order to identify such factors that recruit Hst2 to the phosphorylated Histone H3 tail, we purified FLAG-tagged Hst2 from yeast extracts (Fig. 1a) and analyzed the co-purifying proteins by mass spectrometry. We identified several novel interactors of Hst2 with high fold difference compared to the negative control (untagged Hst2) (Supplementary Fig. S2 and Supplementary Table 3). Gene ontology (GO term) analysis revealed a strong overrepresentation of proteins involved in chromatin related processes, such as replication (Fig. 1b). We confirmed the interaction of Hst2 with Mcm2, a component of the MCM helicase using Western blot (Supplementary Fig. S3). However, since the association of MCM helicase with chromatin is regulated by other mechanisms than H3 phosphorylation (23), this complex is unlikely to recruit Hst2 to phosphorylated H3 tails.

**Figure 1:**
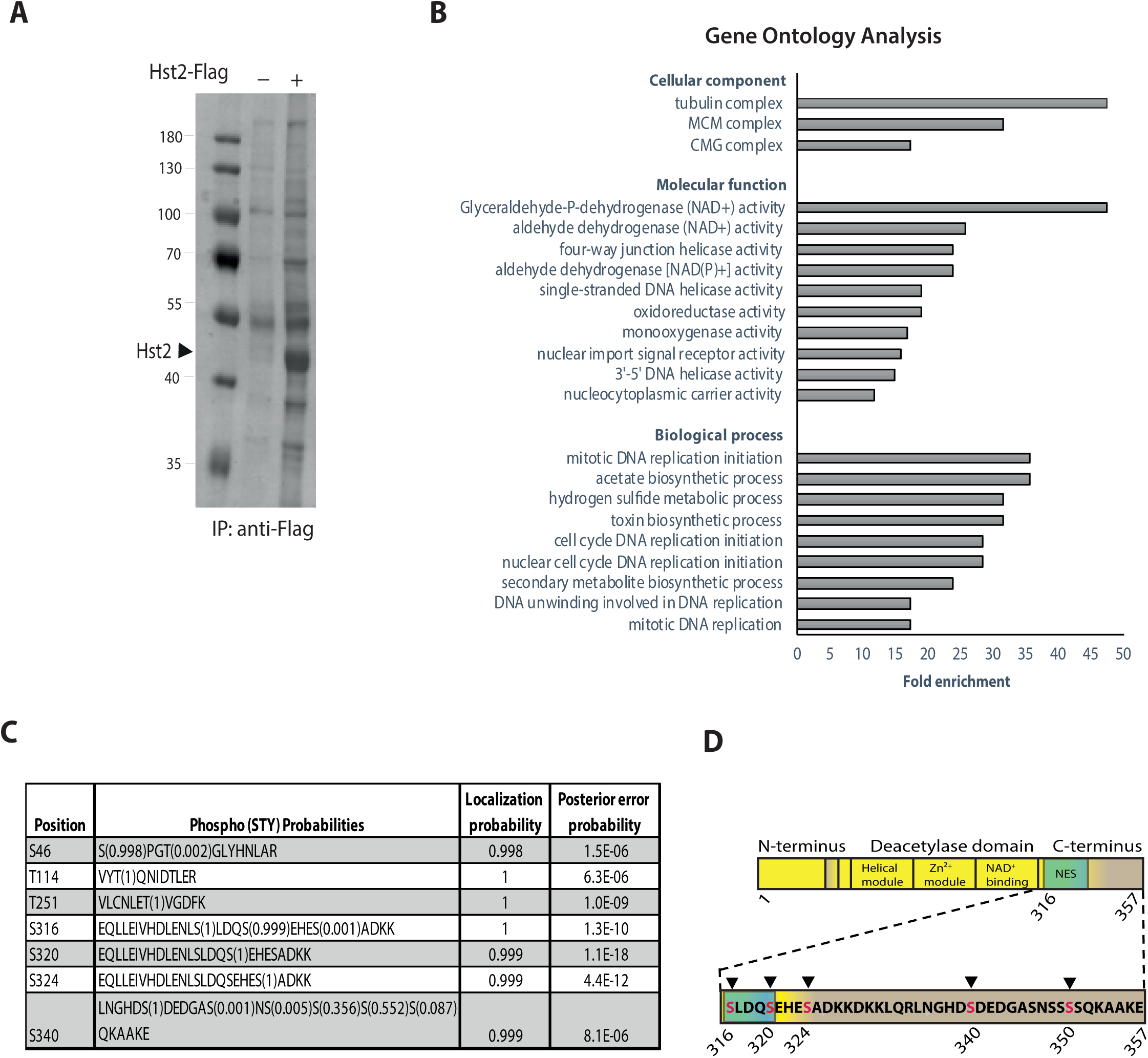
Identification of Hst2 interaction partners and PTMs by mass-spectrometric analysis. (A) Immunoprecipitation of Flag-tagged Hst2. Yeast cells expressing Flag-Hst2 were isolated using ANTI-FLAG^®^ M2 beads, analyzed by 10% SDS-PAGE and stained with Instant Blue. Cells expressing untagged Hst2 served as control. (B) Gene ontology analysis of differentially associated proteins identified in A. Analysis was performed using the online tool “The Gene Ontology Resource”. (C) Phosphorylation sites identified on Hst2 by MS/MS analysis. (D) Schematic representation of Hst2 C-terminal phosphorylation sites. NES: Nuclear Export Sequence (amino acids 306-317).

If the Hst2-H3 interaction requires the presence of PTMs on Hst2, the abundance of the bridging factors in the pull-down might be very low because of the usually low stoichiometry of PTMs. Therefore, we analyzed Hst2 for the presence of phosphorylated residues by MS/MS and identified five serine and two threonine phosphorylation sites (Fig. 1c). Most sites reside in the unstructured C-terminus of the protein. One of the residues, S340, had been reported previously as being phosphorylated (24). Three serine phosphorylation sites (S316, S320 and S324) cluster downstream of the nuclear export sequence (NES) in the C-terminus (Fig. 1d) and may therefore be involved in regulating nuclear export.

We hypothesized that phosphorylation of Hst2 is essential for its recruitment to H3 S10ph. Typical mediators of interactions between two phosphorylated proteins are 14-3-3 proteins (25-27). 14-3-3 proteins are involved in numerous important cellular processes such as transcription (28) and cell cycle control (29) and are consequently associated with diseases including neurodegenerative disorders and cancer (30).

The interaction of 14-3-3 proteins with H3 S10ph is well established across different organisms (31-33). Conversely, several KDACs are known to interact with 14-3-3 proteins in a phosphorylation-dependent manner (34). For example, SirT2 interacts with 14-3-3β/γ in humans (35) and the homologous Sir2.1 and PAR-5/FTT-2 interact in nematodes (36,37). Therefore, we tested whether deletion of the budding yeast 14-3-3 proteins, Bmh1 and Bmh2, interferes with deacetylation of H4 K16 by Hst2 in mitosis, an established hallmark of short-range chromatin compaction (17). Yeast cells lacking BMH1 that were metaphase-blocked with nocodazole indeed showed elevated H4 K16ac levels similar to cells without HST2 (Fig. 2a). Deletion of BMH2 had the same effect to a lesser extent. In line with our assumption that Bmh1 mediates the interaction of phosphorylated Hst2 with H3 S10ph, removing the unstructured C-terminus of Hst2 was sufficient to prevent H4 K16 deacetylation.

**Figure 2:**
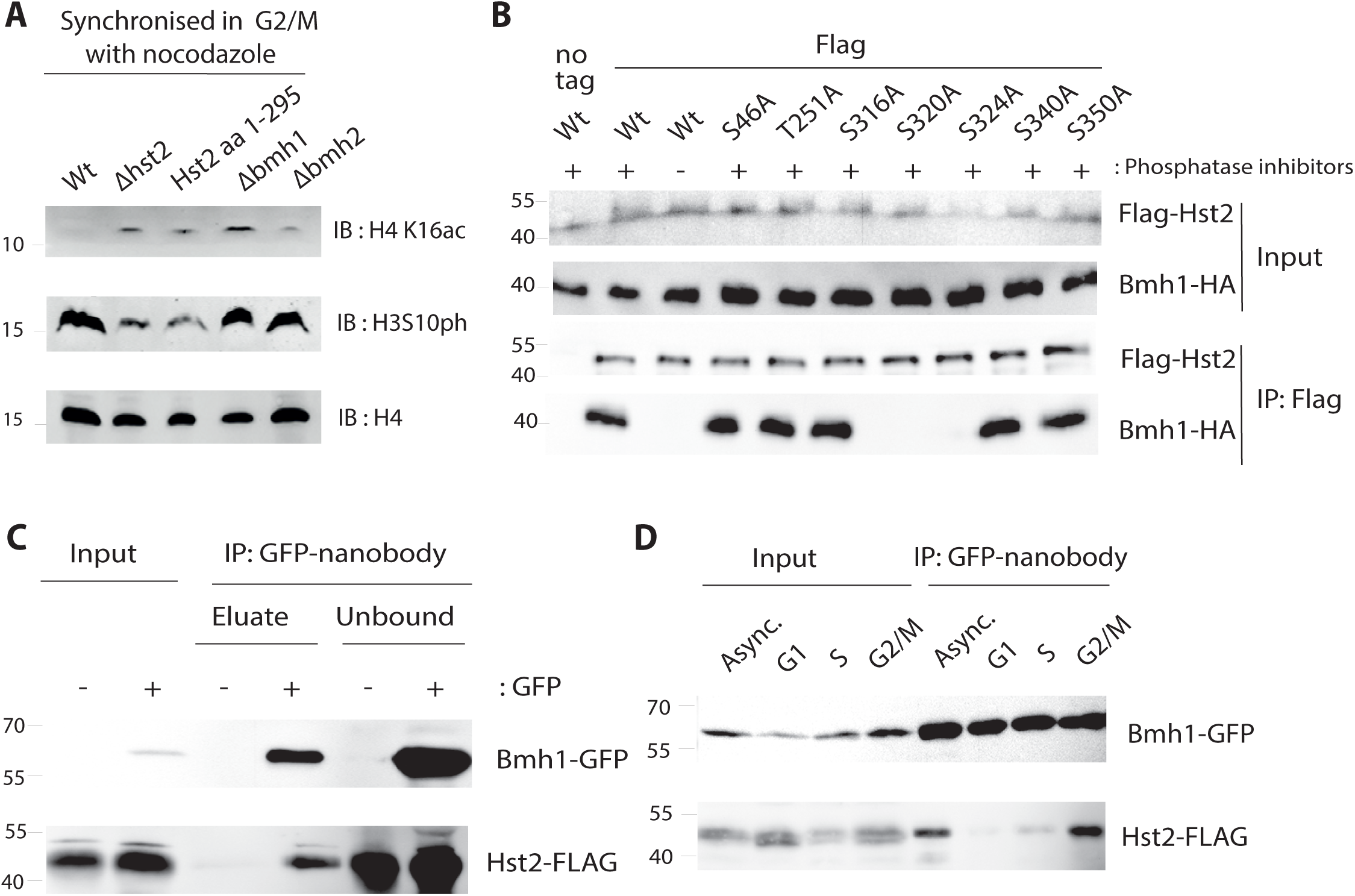
Interaction of Hst2 with 14-3-3 protein Bmh1 is required for H4 K16ac deacetylation in mitosis. (A) Bmh1 and the C-terminus of Hst2 are essential for H4 K16ac deacetylation in mitosis. Yeast cells were arrested in M phase with nocodazole and whole cell lysates analyzed by Western blot with the indicated antibodies. (B) Interaction of Bmh1 with Hst2 *in vivo* depends on phosphorylation of Hst2 S320 or S324. (C) Flag-Hst2 proteins were immuno-purified from overexpressing yeast cells and proteins analyzed by Western blot with the indicated antibodies. (D) Bmh1 interacts with Hst2 *in vivo* at endogenous levels. Bmh1-GFP was isolated from yeast cells with or without genomically Flag-tagged Hst2 using GFP-nanobody beads. Bound proteins were eluted with SUMO protease (releasing the nanobody) and analyzed by Western blot. (E) Bmh1 interacts with Hst2 *in vivo* at endogenous levels only in G2/M phase of mitosis. Yeast cells with genomically Flag-tagged Hst2 and GFP-tagged Bmh1 either asynchronous or arrested with α-factor (G1), Hydroxyurea (S phase) or nocodazole (G2/M), respectively, were lysed and Bmh1-GFP isolated and bound protein analyzed as in C.

Next, we tested whether Bmh1 and Hst2 interact physically. Indeed, when we immune-precipitated Hst2 (overexpressed with N-terminal FLAG-tag), Bmh1 efficiently co-purified only in the presence of phosphatase inhibitors, indicating a phosphorylation-dependent interaction between both proteins (Fig. 2b). The interaction further depended on the presence of serine residues 320 and 324, which we had shown to be phosphorylated in Hst2 (Fig. 1c). Mutating the other phosphorylation sites did not affect the interaction with Bmh1. Interestingly, mutation of either Ser-320 or Ser-324 to alanine alone was sufficient to completely abolish Bmh1 binding. Because optimal sequence motifs for 14-3-3 proteins are R-X_2-3_-(pS/pT)-X-P (38), we consider it unlikely that Bmh1 requires the presence of both phosphorylations for binding. It seems more likely that the two phosphorylation sites interact functionally, for example, that phosphorylation of one site depends on the presence of a serine residue at the other site.

Since overexpression of Hst2 might artificially induce the interaction with Bmh1, we repeated the pulldown experiments at endogenous protein levels. Therefore, we precipitated Bmh1-GFP from yeast lysates with genomically FLAG-tagged Hst2 (Fig. 2c). Hst2-FLAG specifically co-eluted with Bmh1 depending on the GFP-tag on the latter, confirming that the Hst2-Bmh1 interaction also occurs at endogenous protein concentrations.

Next, we asked whether the Hst2-Bmh1 interaction is influenced by the cell cycle stage. Therefore, we repeated the pulldown experiments at endogenous protein levels with cell cycle synchronized yeast cultures (Fig. 2d). Pulldown experiments with lysates from yeasts blocked in G1-phase with α-factor or S-phase with hydroxyurea showed very little interaction of Hst2 with Bmh1. In contrast, we observed efficient interaction of Hst2 with Bmh1 when cells were blocked in metaphase with nocodazole. This result agrees with our model that Bmh1 recruits Hst2 to chromatin in mitosis when it is needed for short-range chromosome compaction.

To test the Hst2-Bmh1 interaction under defined conditions, we prepared singly phosphorylated Hst2 proteins in *E. coli* using genetic code expansion (20). Thereby, phosphoserine is encoded in response to amber (UAG) stop codons by an archaeal phosphoseryl-tRNA synthetase/tRNA_CUA_ pair introduced in an *E. coli* strain lacking phosphoserine phosphatase SerB. Suppression of amber codons replacing the codon for phosphorylated serine residues in Hst2 results in the production of site-specifically phosphorylated protein. We produced Hst2 S320ph and Hst2 S324ph and compared their ability to bind to Strep-tagged Bmh1 with unphosphorylated Hst2 (Fig. 3a). Both phosphorylated forms of Hst2 were efficiently recovered in these pulldown experiments, while unmodified Hst2 did not associate with Bmh1. Hence, both serine residues mutually required for the Hst2-Bmh1 interaction *in vivo* are individually sufficient to mediate the interaction *in vitro*.

**Figure 3:**
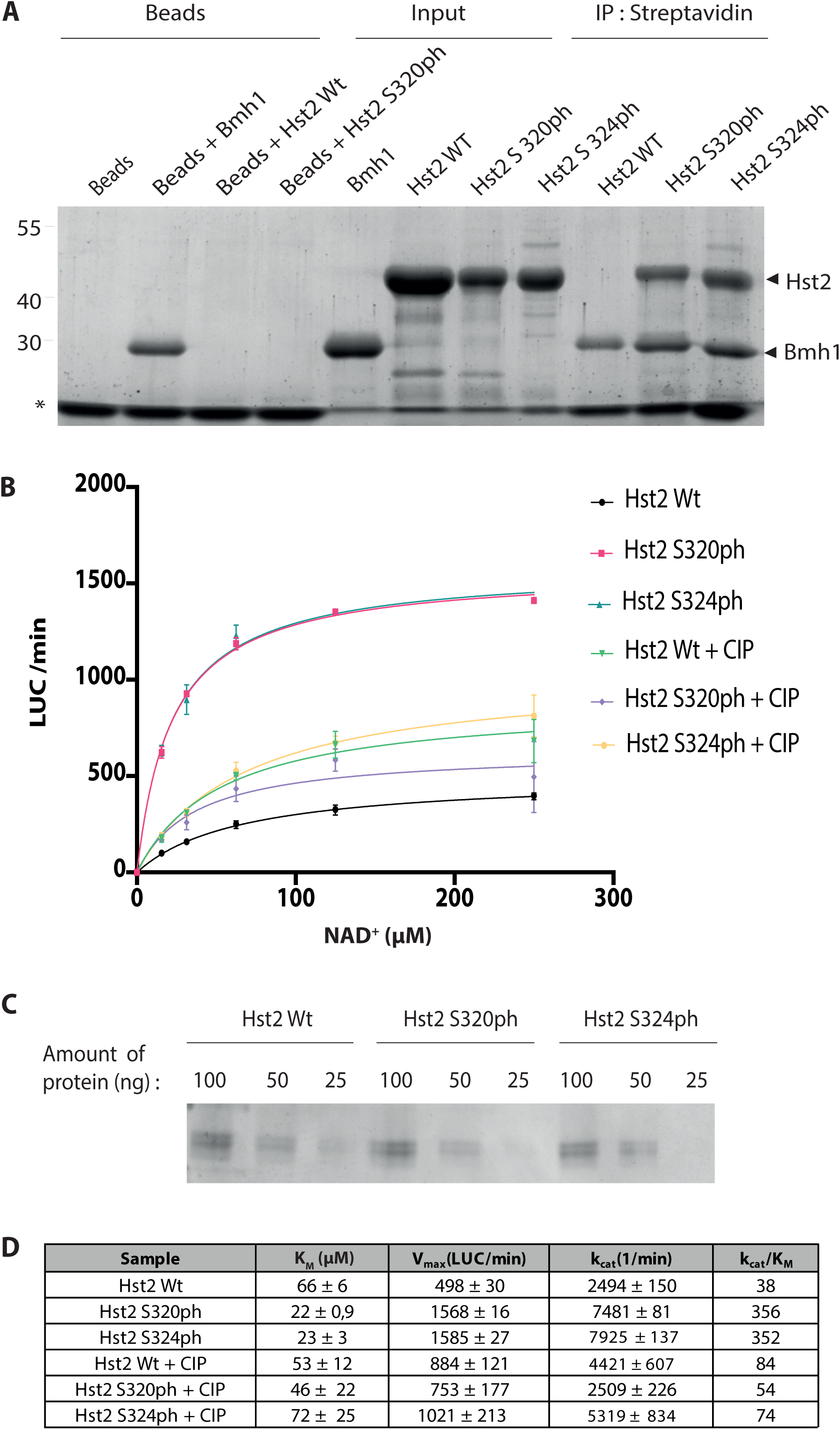
Phosphorylation of Hst2 enhances catalytic activity and mediates interaction with Bmh1 *in vitro.* (A) Bmh1 binds to singly phosphorylated Hst2 *in vitro*. Phospho-forms of Hst2 produced in *E. coli* were incubated with Strep-tagged Bmh1 on streptavidin beads. Bound proteins were eluted with SDS sample buffer, analyzed by 10% SDS-PAGE and stained with Instant Blue. * denotes streptavidin bead specific band. (B) Kinetic analysis of Hst2 and the phospho-forms S320ph and S324ph. Deacetylation of FLuc K529ac by Hst2 isoforms was measured in continuous assay format. CIP was used to remove phosphorylation of Hst2. (C) Equal amounts of Hst2 proteins analyzed by 10% SDS-PAGE as loading control for B. (D) Kinetic parameters of Hst2 and phospho-Hst2. Error values are calculated using a root mean least squares approach from a double-reciprocal plot.

The phosphorylation of Hst2 may have additional functions in the regulation of Hst2 activity. For example, phosphorylation of S316 could regulate nuclear export of Hst2 because this residue is part of the NES. Furthermore, the C-terminal α-helix of Hst2 has been shown to interfere with NAD^+^-binding (39). Therefore, we measured the activity of recombinant Hst2 with or without phosphorylation of S320 or S324 in dependence of NAD^+^-concentration using acetylated Firefly luciferase as substrate (21) (Fig. 3b). The phosphorylated forms of Hst2 both showed a threefold increase in Vmax and a threefold decrease in K_M_ for NAD^+^, resulting in an almost tenfold higher catalytic efficiency. The enhanced catalytic efficiency was reverted by preincubation of the phosphorylated proteins with calf intestine phosphatase (CIP), confirming that the presence of the phosphorylation is essential for the effect.

This indicates that the unphosphorylated Hst2 C-terminus acts like a mixed noncompetitive inhibitor that is masked by phosphorylation. This can be interpreted in the way that binding of the C-terminus to the catalytic core reduces the affinity for NAD^+^ (increasing K_M_) and also slows catalysis (reducing Vmax). The regulation of catalytic activity by phosphorylation of C-terminal residues appears to be a conserved mechanism that has also been observed for the human Hst2 homologue, SirT2 (40,41).

## Conclusions

Chromosome condensation in mitosis is licensed by kinetochores via the recruitment of shugoshin and Hst2 (19). Phosphorylation of H3 S10 by Aurora kinase B is a central event in this process that mediates deacetylation of H4 K16ac by Hst2 (17,22). Here, we demonstrate that binding of Hst2 to this mark is mediated by Bmh1, which simultaneously binds the phosphorylated H3 tail and C-terminally phosphorylated Hst2 (Fig. 4).

**Figure 4:**
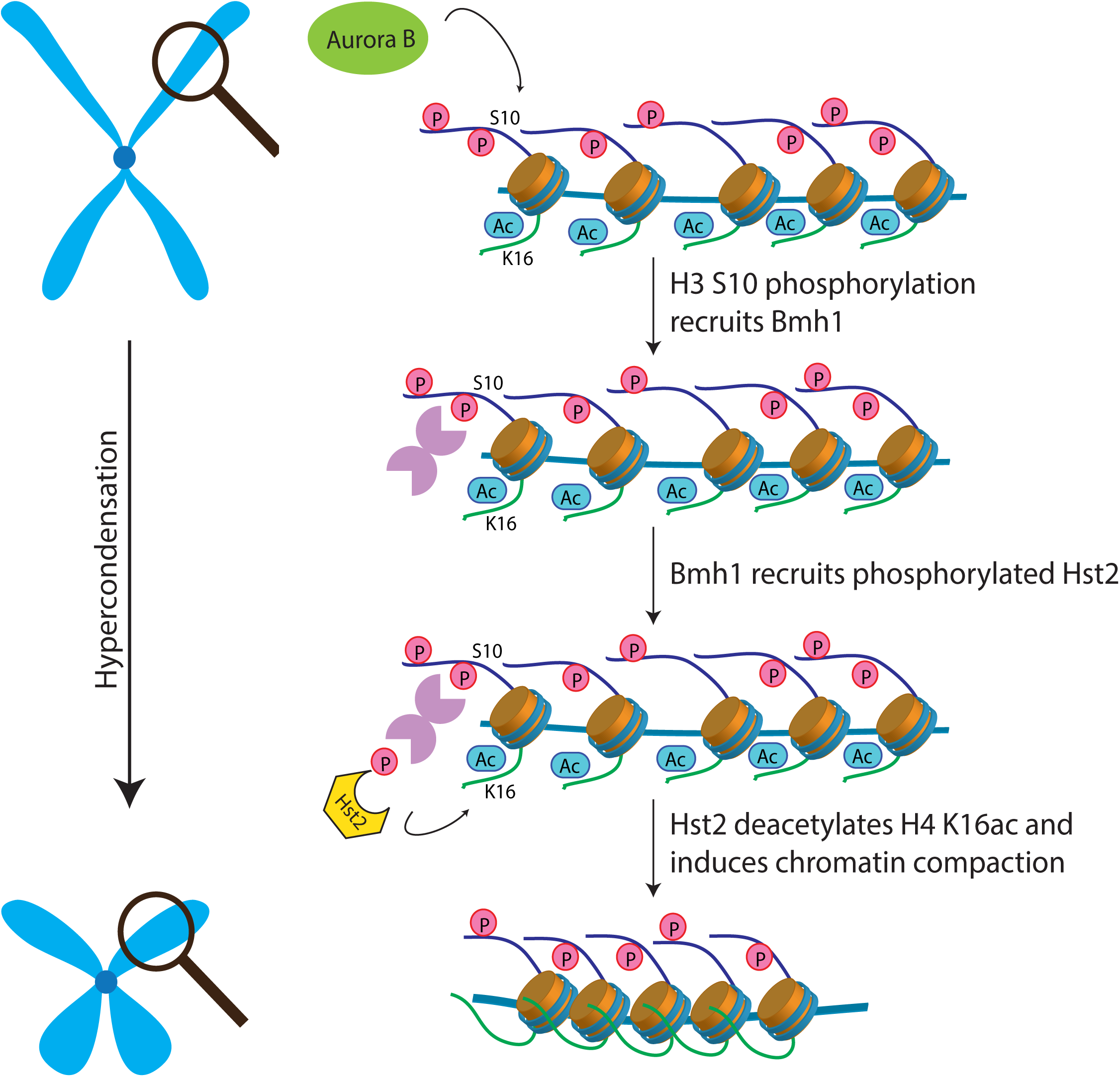
Bmh1-mediated recruitment of phosphorylated Hst2 to mitotic chromatin induces chromatin compaction.

Our results show that phosphorylation of Hst2 S320 or S324 is necessary and sufficient to induce the interaction with Bmh1. In contrast, mutation of either site to alanine is sufficient to prevent co-immunoprecipitation of Hst2 and Bmh1 *in vivo*. This may be the result of a complex interplay between these sites *in vivo* such that the phosphorylation of one site depends on the presence of a serine residue at the other site. Alternatively, the *in vitro* pulldown experiments may require less affine interactions because the interaction partners are present at much higher concentration in the assay. Interestingly, recruitment of human SirT2 to chromatin is also regulated by phosphorylation (40,42), suggesting that a similar mechanism might act in mammalian cells.

An important open question is how compaction spreads from kinetochores along chromosome arms. The multi-valent nature of the interaction of chromatin, Bmh1-dimers and Hst2 (which can form trimers *in vitro* (39)) may facilitate liquid-liquid phase separation. This hypothesis is supported by the presence of poly-glutamine stretches at the C-terminus of Bmh1 and Bmh2. These motifs are known to interact with nucleic acids and are often involved in phase-separation processes (43).

## Acknowledgements

We thank Petra Geue for technical assistance and all members of the Neumann group for helpful discussions.

## Author Contributions

Conceptualization, H.N.; Methodology, N.J., P.J., Investigation, N.J., P.J., H.N.; Data Curation, P.J.; Writing – Original Draft, N.J., H.N.; Writing – Review & Editing, N.J., P.J., H.N.; Supervision, H.N.; Project Administration H.N.; Funding Acquisition, H.N.

